# *Thermosynechococcus* switches the direction of phototaxis by a c-di-GMP dependent process with high spatial resolution

**DOI:** 10.1101/2021.08.26.457869

**Authors:** Daisuke Nakane, Gen Enomoto, Annegret Wilde, Takayuki Nishizaka

## Abstract

Many cyanobacteria, which use light as an energy source via photosynthesis, show directional movement towards or away from a light source. However, the molecular and cell biological mechanisms for switching the direction of movement remain unclear. Here, we visualized type IV pilus-dependent cell movement in the rod-shaped thermophilic cyanobacterium *Thermosynechococcus vulcanus* using optical microscopy at physiological temperature and light conditions. Positive and negative phototaxis were controlled on a short time scale of 1 min. The cells smoothly moved over solid surfaces towards green light, but the direction was switched to backward movement when we applied additional blue light illumination. The switching was mediated by three photoreceptors, SesA, SesB and SesC, which have cyanobacteriochrome photosensory domains and synthesis/degradation activity of the bacterial second messenger cyclic dimeric GMP (c-di-GMP). Our results suggest that the decision-making process for directional switching in phototaxis involves light-dependent changes in the cellular concentration of c-di-GMP. Furthermore, we reveal that rod-shaped cells can move perpendicular to the light vector, indicating that the polarity can be controlled not only by pole-to-pole regulation but also within-a-pole regulation. This study provides insights into previously undescribed rapid bacterial polarity regulation via second messenger signalling with high spatial resolution.

## Introduction

Cyanobacteria are phototrophic microorganisms, and optimal light conditions are crucial for efficient photosynthesis. Therefore, several cyanobacterial strains are able to move either into their preferred light habitat or away from a harmful or stressful environment (1). This decision-making process is based on sensing the light direction and light intensity and quality on a short time scale. Moderate red light is used as the preferred energy source for oxygenic photosynthesis, while strong light or UV light causes cell damage (2). However, how cyanobacterial cells rapidly switch the direction of movement remains unclear.

Cyanobacterial movement is usually driven by type IV pili (T4P) (1), a general bacterial molecular machine. This machine enables cellular movement by repeated cycles of extension and retraction of the pili (3, 4). T4P are often localized at the cell poles in rod-shaped bacteria, and their localization at a certain pole is dynamically controlled to achieve directional movement (5). Bacteria, such as *Pseudomonas*, exhibit chemotactic behaviour to activate T4P at the leading pole on a time scale of hours. Consequently, the longer axis of cells is roughly aligned in parallel along the chemical gradient (6). Cyanobacterial cells make a decision of directional movement in a few minutes upon light sensing by dedicated photoreceptors (7). Once the lateral light stimulus is applied to the cells, they detect the orientation of a light source using their own cell body as an optical lens. In the coccoid *Synechocystis* sp. PCC 6803 (*Synechocystis*), the light is focused at the cell envelope, thereby generating a light spot opposite to the light source. Such microoptic effects have been described in cyanobacteria of different shapes (7, 8). Yang *et al*. suggested that the rod-shaped cyanobacterium *Synechococcus elongatus* UTEX 3055 also utilizes the micro-optics effect to sense directional light by the polar-localized photoreceptor PixJ (9). However, T4P-dependent cell behaviour has not been clarified at the single-cell level in rod-shaped cyanobacteria (9). Furthermore, since most of the knowledge of phototaxis is derived from coccoid-shaped *Synechocystis*, how the polarity of the cell structure is involved in phototaxis regulation is not clear.

The nucleotide second messenger molecule c-di-GMP is the critical molecule governing bacterial motility as the master regulator of bacterial lifestyle transitions (10, 11). The high intracellular concentration of c-di-GMP is universally implicated in the repression of motility and induction of sessile multicellular community development. A decrease in c-di-GMP levels is often required for optimum motile behaviour involving type IV pili or flagella (10). The diguanylate cyclase (DGC) activity mediated by GGDEF domains is necessary to produce c-di-GMP, whereas EAL and HD-GYP domains harbour phosphodiesterase (PDE) activities to degrade c-di-GMP. These domains are often combined to signal sensory domains (12), enabling bacterial cells to integrate multiple environmental information into cellular c-di-GMP levels to orchestrate various cellular responses to accomplish complex lifestyle transitions.

In this study, we establish a microscopy setup to analyse the phototaxis of the rod-shaped thermophilic cyanobacterium *Thermosynechococcus vulcanus* using live cells at the single-cell level. We dissect the contributions of different light colours and evaluate the functions of each putative photoreceptor gene. We show that both positive and negative phototaxis in *T. vulcanus* were controlled by a specific green-to-blue light ratio. Furthermore, we provide evidence that the reversion of phototaxis is mediated by photoreceptors and their activity in the synthesis or degradation of cyclic di-GMP. The asymmetric distribution of T4P at both cell poles achieves directional movement with a random orientation of the cells along their optical axis. We suggest a within-a-pole regulation of polarity that governs the directional movement of *T. vulcanus* and requires signalling at high spatiotemporal resolution. The light dependency of the phototactic response is consistent with a proposed model of *T. vulcanus* behaviour in a mat, which is their natural habitat.

## Results

### Control of positive and negative phototaxis

*T. vulcanus* used in this study was originally derived from the strain at the culture collection of NIES-2134 (http://mcc.nies.go.jp/). Since the original strain exhibited a heterogeneous phenotype of phototaxis under optimal growth conditions, a clone showing clear positive phototaxis in moderate light was isolated (13), and its complete genome was sequenced (AP018202). This clone was used as wild type (WT) in the following experiments. The strain also showed negative phototaxis when we applied a strong light stimulus. This bidirectional phototaxis was visualized as colony migration on a BG11 agar plate in a long-term observation for 8 hours (Fig. 1A), consistent with previous data based on a closely related strain (14).

**Fig. 1.**
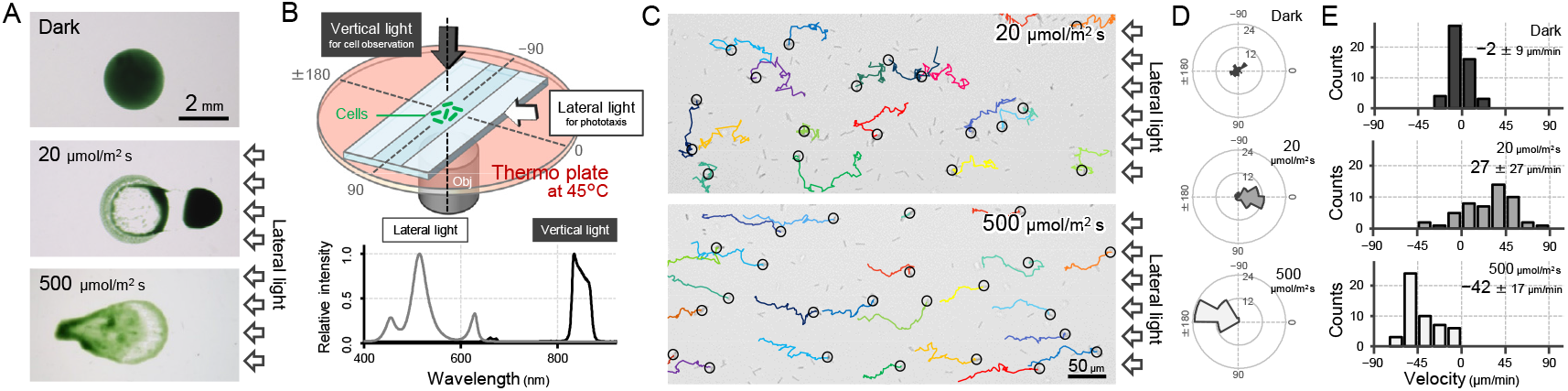
Positive and negative phototaxis of *T. vulcanus*. (A) Phototaxis on agar plates. Images were taken 8 hours after spotting the cell suspension. (B) Diagram of the experimental setup to visualize single cellular behaviour under optical microscopy. The glass chamber was heated at 45 °C with a thermoplate on a microscope stage. Vertical and lateral light sources were used for cell observation and stimulation of phototaxis, respectively. Light spectra are presented at the bottom. (C) Bright-field cell image and their moving trajectories for 120 s (colour lines) on a glass surface. The cell at the start position of a trajectory is marked by the black circle. The white arrows on the right side of the image represent the direction of the light axis. (D) Rose plots under dark, weak light, and strong light illumination. The moving direction of a cell that translocated more than 6 μm min^-1^ was counted. Angle 0 was the direction towards the lateral light source (N = 50 cells). (E) Histograms of the cell velocity along the lateral light axis. The cell displacement for a duration of 1 minute was measured at 4 min after lateral illumination was turned on (N = 50 cells). Cell movements towards the light source are shown as a positive value.

To observe a single-cell trajectory during phototaxis on a short time scale, we constructed an optical setup that allowed us to stimulate the cell with lateral light on a microscope stage, which was heated at a temperature of 45 °C to observe the physiological cell behaviour (Fig. 1BC). The position of the rod-shaped cell was visualized by near-infrared light through a bandpass filter with a centre wavelength of 850 nm and a full width at half maximum of 40 nm from a halogen lamp with a fluence rate of 1 μmol m^-2^ s^-1^, which was confirmed to have no effect on motility (Fig. 1DE) (7, 14). When we applied a lateral white illumination of 20 μmol m^-2^ s^-1^, the cells started to exhibit directional movement towards the light source in a few minutes (Fig. 1C *top*). On the other hand, the cells showed directional movement away from the light source upon strong white light illumination of 500 μmol m^-2^ s^-1^ (Fig. 1C *bottom*). The rose plot, a round histogram that simultaneously presents the number of occurrences and direction, indicates that lateral light illumination induced both positive and negative phototaxis (Fig. 1D). The velocity of cell movement along the optical axis on a glass surface was calculated to be 20–50 μm min^-1^ (Fig. 1E), which is 5–10 times faster than that of *Synechocystis* or *S. elongatus* strain UTEX 3055 (7, 9). These data clearly show that the positive and negative phototaxis is controllable on this microscopic setup. Therefore, we started with a more detailed quantitative analysis of the cell movement of *T. vulcanus*.

### Wavelength dependence of phototaxis

We set up 5 LEDs with various wavelengths as lateral light sources to stimulate the phototaxis of cells under the microscope (Fig. 2A). We used LEDs for blue, teal, green, red, and far-red light with peak wavelengths at 450, 490, 530, 625, and 730 nm, respectively, and spectral bandwidths in a range of 15–40 nm (Fig. 2B). When we applied monochromatic illumination with green or teal light at a rate of 70 μmol m^-2^ s^-1^, the cells showed directional movement within a few minutes (Fig. 2C and Movie S1). The cell velocity along the optical axis was 30 μm min^-1^ in the green and 20 μm min^-1^ in the teal (Fig. S1). Monochromatic illumination with blue or red light induced the fast formation of small aggregates that moved randomly (Fig. 2C, Fig. S1 and Movie S2). The previously described macroscopic floc formation of *T. vulcanus*, which involves the induction of cellulose synthesis (15), is a long-term response that is referred to as cell aggregation. For clarity, in the current study, we call the small aggregates observed under the microscope, which are formed very quickly, microcolonies.

**Fig. 2.**
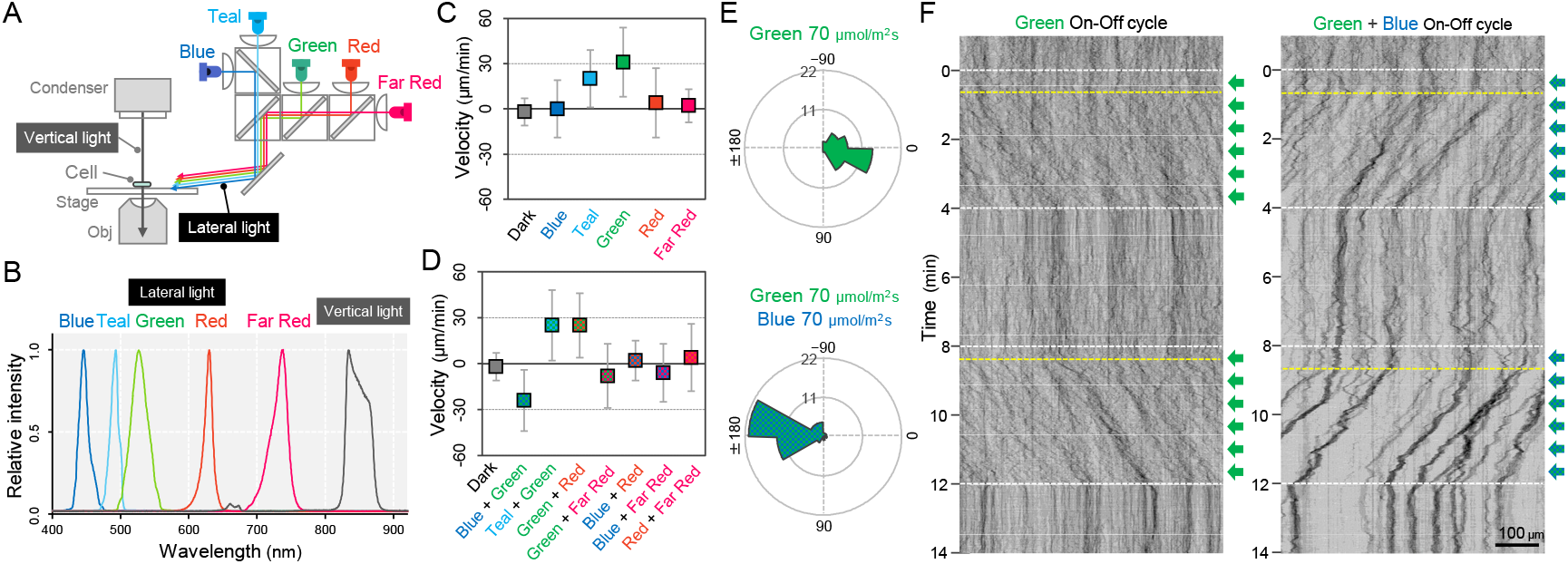
Wavelength dependency of phototaxis via optical microscopy. (A) Schematics of the lateral illumination for phototaxis. Five LEDs were simultaneously applied through dichroic mirrors from the right side. (B) Spectra of lateral and vertical light for phototaxis. (C) Effects of the monochromatic light source on the phototactic behaviour of cells on a glass surface. (D) Effects of dichromatic light source on the phototactic behaviour of cells over the glass surface. Each lateral light was used at a fluence rate of 70 μmol m^-2^ s^-1^. The average and standard deviation (SD) of the cell velocity along the light axis are presented (N = 50). (E) Rose plots under green light at 70 μmol m^-2^ s^-1^ (upper) and green and blue light at 70 μmol m^-2^ s^-1^ (lower). The moving direction of a cell that translocated more than 6 μm min^-1^ was counted. Angle 0 was the direction towards the lateral light source (N = 50 cells). (F) On-off control of phototaxis. A kymogram of cell movements along the optical axis of lateral illumination is presented. Directional movements of cells are shown by the tilted lines over time. The tilted lines from the left-upper to the right-lower side and from the right-upper to the left-lower side presented positive and negative phototaxis, respectively. Lateral illumination was applied with a time interval of 4 min and indicated by the dashed white lines (see also Movie S5 and Movie S6). The delay of the cell response after the illumination was turned on is indicated by the dashed yellow lines.

In summary, our data suggest that phototaxis in *T. vulcanus* has a wavelength dependence on green light for positive phototaxis. The cell velocity was increased from 20 to 70 μmol m^-2^ s^-1^ of green light and remained positive even at 700 μmol m^-2^ s^-1^, which is roughly one-third of the intensity of direct sunshine (Fig. S2). Note that under nonphysiological conditions at room temperature (25 °C), the cells showed negative phototaxis under lateral green light due to a so far unknown mechanism (Fig. S3 and Movie S3). Therefore, we performed all further experiments at the normal growth temperature of *T. vulcanus* at 45 °C using the special microscopy setup developed here (see Materials and Methods section).

Next, we observed cell movement with a combination of two LEDs of different wavelengths simultaneously. We found that the direction of the phototactic response was reversed after a combination of green and blue light (Fig. 2D and Movie S4), whereas the cells retained positive movement when green was combined with teal or red light. No directional movement was observed without green light (Fig. S4). These data suggest that blue light is responsible for the directional switch of phototaxis. To study these effects in more detail, we applied constant green light intensity at 70 μmol m^-2^ s^-1^ and combined it with different blue light intensities (Fig. S5). Illumination with 20 μmol m^-2^ s^-1^ blue light in addition to green light led only to a decrease in cell velocity, whereas 200 μmol m^-2^ s^-1^ blue light induced negative phototaxis in all cells. Notably, negative phototaxis was induced even when we applied blue light from the opposite side of the lateral green light source, suggesting that the reversal of the phototaxis from positive to negative does not depend on the direction of blue light illumination (Fig. S6). In the following experiments, we used green light at 70 μmol m^-2^ s^-1^ for positive phototaxis and the combination of green light at 70 μmol m^-2^ s^-1^ and blue light at 200 μmol m^-2^ s^-1^ for negative phototaxis (Fig. 2E). Under these conditions, the kymographs representing the spatial position of the cells along the lateral light axis over time depict examples of both positive and negative phototaxis as tilted lines (Fig. 2F). Directed movement was barely observed for the first 30 sec after the light was turned on in most cells under repeated on-off cycles of lateral illumination (Movie S5 and Movie S6). This delay has also been reported in phototaxis of *Synechocystis* (7).

### Photoreceptors for phototaxis

To identify the photoreceptors involved in the phototaxis of *T. vulcanus*, we constructed mutants lacking the genes for the cyanobacteriochromes SesA (16), SesB and SesC (17), the putative circadian input protein CikA (18), the BLUF-domain protein PixD (19), the putative photoreceptor LOV (20), and the orange-carotenoid protein OCP (21), all of which are predicted to sense within the blue-to-green light region (Fig. 3A). Note that no red/far-red absorbing phytochrome photoreceptor gene has been identified in the complete genome of *T. vulcanus*. All mutants showed positive phototaxis towards green light (Fig. S3B, upper), suggesting that none of these photoreceptors is involved in the control of positive phototaxis. Next, we applied green and blue light to check negative phototaxis and measured the velocity of cell displacement along the light axis (Fig. 3B, lower). Whereas WT cells showed negative phototaxis upon lateral illumination with green and blue light, the *ΔsesA* mutant maintained positive phototaxis. SesA harbours a GGDEF domain and synthesizes the second messenger c-di-GMP in response to blue light (17). This suggests that the directional switch from positive to negative phototaxis may be triggered by SesA-dependent c-di-GMP synthesis. This assumption is supported by measurements of the intracellular c-di-GMP concentration (Fig. S7). The WT showed a more than three times higher c-di-GMP content under blue light than under the other tested light conditions, such as green, red, and white light. In contrast, *ΔsesA* lost the blue light-dependent increase in intracellular c-di-GMP levels.

**Fig. 3.**
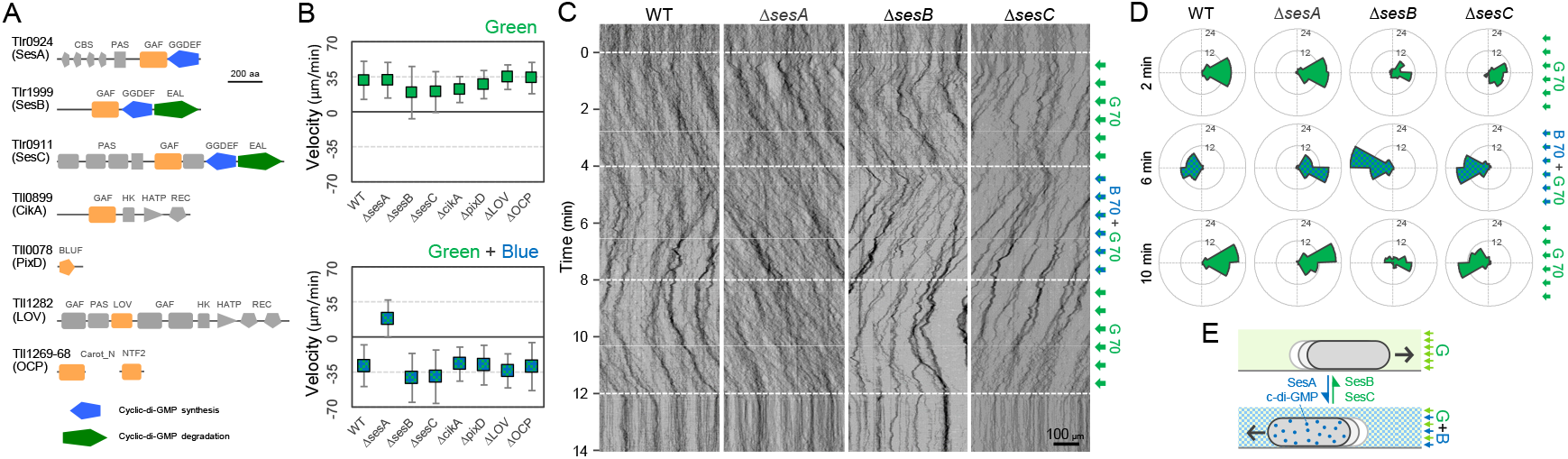
Photoreceptors for phototaxis. (A) Domain composition of candidate photoreceptor-containing proteins in *T. vulcanus*. (B) Mutant cell behaviour after lateral light illumination. *Upper*: Lateral green light at a fluence rate of 70 μmol m^-2^ s^-1^. *Lower*: Lateral green and blue light at a fluence rate of 70 μmol m^-2^ s^-1^. The cell displacement for a duration of 1 minute was measured at 3 min after lateral illumination was turned on (N = 50 cells). Cell movements towards the light source are shown as a positive value. (C) Kymograph of cell movements in the WT, Δ*sesA*, Δ*sesB*, and Δ*sesC* mutants along the optical axis of lateral illumination. The cell position was visualized by near-infrared light. Phototaxis was stimulated by the lateral illumination of green and green/blue light. Green light was applied from time 0 to 4 and from 8 to 12 min, and green/blue light was applied from time 4 to 8 min (see also Movies S7-S10). (D) Rose plot. The moving direction of a cell that translocated more than 6 μm min^-1^ was counted. Lateral light was applied from the right side. The cell displacement was measured at each time point, as presented on the left (N = 50 cells). The data come from Panel C. (E) Schematic model of the reversal in phototaxis induced by the intracellular concentration of cyclic di-GMP.

Previous reports have shown that the three photoreceptor proteins SesA, SesB, and SesC work together for c-di-GMP-dependent control of cell aggregation via the regulation of cellulose synthesis (16, 17). SesB degrades c-di-GMP, and its activity is upregulated under teal light irradiation. SesC is a bifunctional protein with enhanced c-di-GMP-producing activity under blue light and enhanced c-di-GMP-degrading activity under green light. To examine the directional switch of phototaxis in detail, we applied lateral illumination in three phases in the order of green, green/blue, and green with a time interval of 4 min (Fig. 3CD and Movie S7). WT cells clearly showed directional movement and switched from positive to negative and back to positive phototaxis in response to the applied light regime. However, Δ*sesA* mutant cells maintained positive phototaxis under all conditions, even after green/blue illumination (Fig. 3CD and Movie S8). In contrast, the Δ*sesC* mutant switched to negative phototaxis but did not move again towards the light source upon illumination with only green light (Fig. 3CD and Movie S9). The Δ*sesB* mutant switched movement from positive to negative and from negative to positive phototaxis, similar to the WT. However, the Δ*sesB* cells continued to aggregate after the light stimulus had stopped (Fig. 3CD and Movie S10), while the WT dispersed in a few minutes. We hypothesize that negative phototaxis is induced by SesA-dependent c-di-GMP synthesis under blue light, whereas SesC and SesB degrade c-di-GMP under green light, thereby inducing positive phototaxis and controlling the dispersion of microcolonies or cell aggregation (22) (Fig. 3E).

### Components of T4P machinery for phototaxis

T4P is known to be essential for phototactic motility in other cyanobacteria (23–25). To evaluate the contribution of T4P to *T. vulcanus* phototaxis, we tested the phenotype of Δ*pilA, ΔpilB, ΔpilT*, and Δ*hfq* mutants (Fig. 4A). Electron microscopy revealed that the Δ*pilB* and Δ*hfq* mutant cells lacked pilus filaments, whereas WT cells had dozens of filaments localized at each cell pole (Fig. 4BC). As expected, the mutants did not show phototaxis upon green light illumination under the current microscope setup (Fig. 4D). These data showed that *T. vulcanus* has a bipolar localization of T4P, which is necessary to generate directional movement in response to light illumination.

**Fig. 4.**
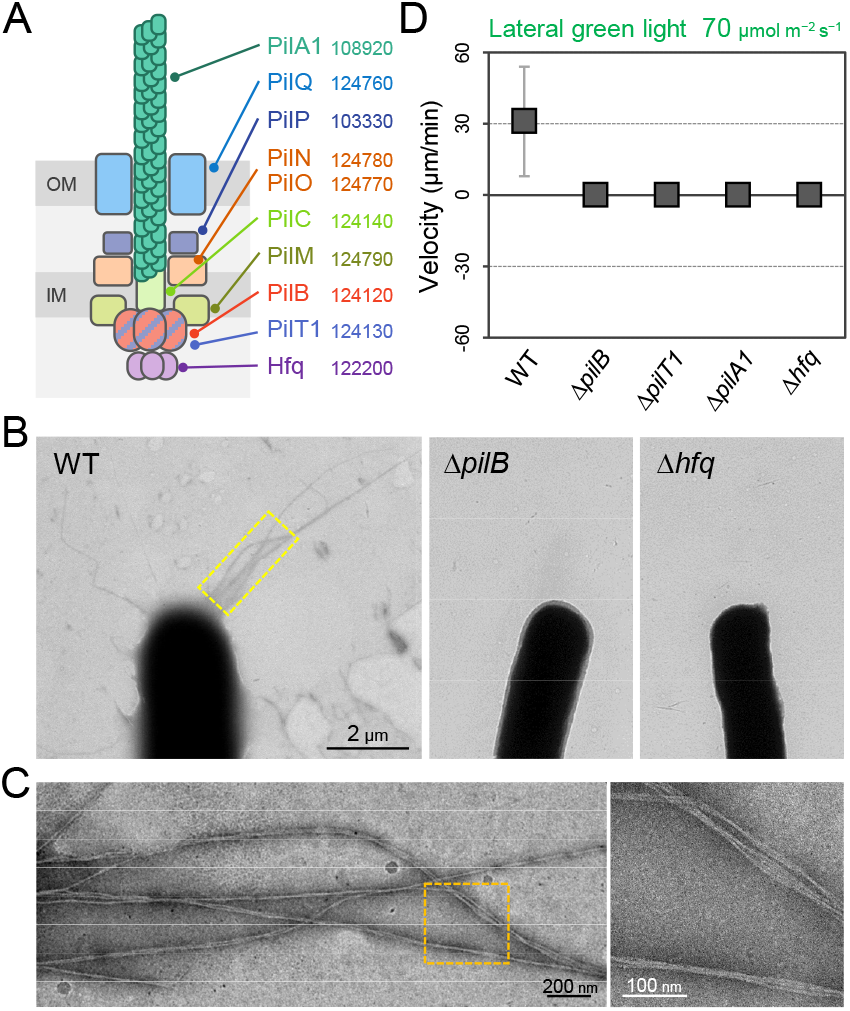
T4P machinery and phototaxis. (A) Schematic of the T4P machinery. Protein components and their gene IDs from the genome of *T. vulcanus* are indicated. The location of the components is presented in reference to other species (3). (B) EM image of a cell. (C) Magnified images of the yellow and orange boxed areas are presented. (D) Effect of lateral light illumination on mutant cells. Lateral green light was applied at a fluence rate of 70 μmol m^-2^ s^-1^. The average and SD of the cell velocity on the glass surface along the light axis are presented (N = 50). Hfq was previously suggested to be involved in the formation of T4P machinery in other cyanobacteria (24).

### Random orientation of cells during negative phototaxis

We observed that cells formed more microcolonies during negative phototaxis than during positive phototaxis (Movie S1 and Movie S4). The cells started to aggregate 1 min after the green and blue light stimulus (Movie S4) but nevertheless kept the directed movement away from the light source (Fig. 5A, upper and Movie S11). Thus, microcolony formation of cells did not hinder directed movements. Under these conditions, approximately 70% of all cells formed such microcolonies and moved directionally, while 30% of cells stayed as single cells (Fig. 5B). These cells were able to adopt a random orientation along the light vector, meaning that they aligned their axis at different angles in relation to the light direction and exhibited directed movements as single cells. As an example, we show one cell aligned perpendicular to the optical axis of lateral light and attached at both poles to the surface (Fig. 5A, left bottom and Movie S12). Another example is a cell positioned in an upright position at one pole (Fig. 5A, right bottom and Movie S13), reminiscent of walking observed previously in *Pseudomonas* (26). Not all single cells aligned in parallel along the light axis (Fig. 5C), suggesting that the force for cell movement is not generated exclusively along the longer axis of the cell.

**Fig. 5.**
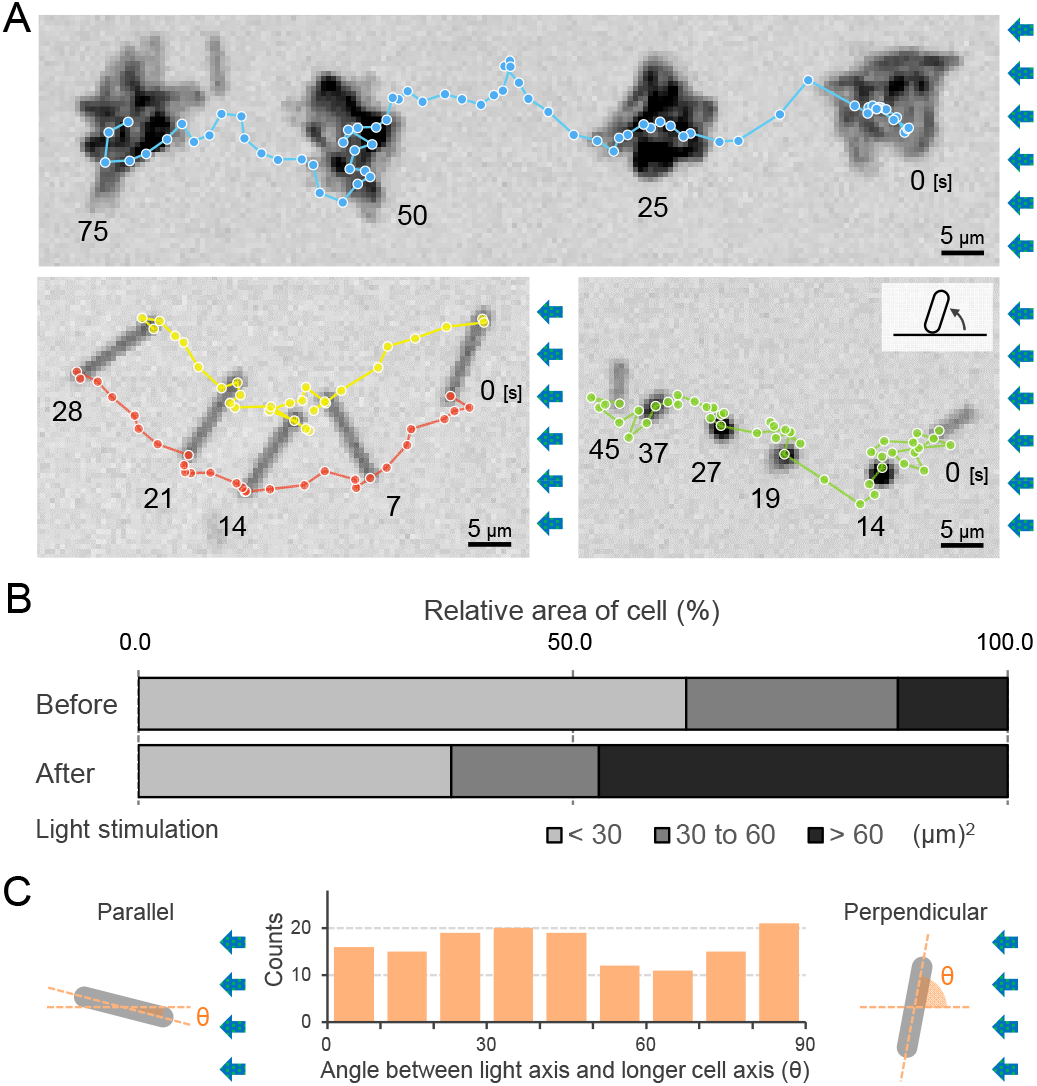
Moving trajectories of cells during negative phototaxis. (A) Cell images and moving trajectories. Images were integrated with a single image at each time duration presented (see also Movies S12-S14). Upper: movement of microcolonies. Left bottom: cell perpendicular to lateral light axis. Right bottom: cell that stood up and kept binding at a cell pole. (B) Cell-cell interaction. The area of a cell moving as a single unit was measured before and after the induction of negative phototaxis and presented as the ratio by the area (N = 100). (C) Moving direction of a cell in relation to the light source. The orientation of single cells during negative phototaxis was measured, and the distribution is presented (N = 100). The absolute angle of the longer axis of a cell was measured in relation to the lateral light axis. The parallel cellular orientation to the light axis (shown on the left) was taken as 0 degrees, whereas the perpendicular orientation (shown on the right) was ideally 90 degrees.

### Direct visualization of T4P dynamics using fluorescent beads

We visualized the T4P dynamics of *T. vulcanus* through beads attached to the pilus fibre (Fig. 6A and Movies S14), as previously described in a coccoid-shaped unicellular cyanobacterium (7). In the presence of 200 nm diameter polystyrene sulfate beads, the beads around the cell pole showed directional movements (Fig. 6BC). The distance between beads and the cell pole increased or decreased with time (Fig. 6D). The bead movement was biased to the cell poles, resulting in the accumulation of beads at the poles after a few minutes of observation (Fig. S8A). This accumulation was not observed in the Δ*pilB* mutant, suggesting that directed bead movement is driven by dynamic T4P activity. The average velocity of beads towards and away from cell poles was measured to be 1.59 ± 0.34 and 2.68 ± 0.33 μm s^-1^ (Fig. 6E), corresponding to the retraction and extension of T4P, respectively (7). The distribution of the velocity was not changed during illumination with green and green/blue light (Fig. S9). Note that the sulfate beads were effectively captured by T4P, whereas neither carboxylate-nor amine-modified beads accumulated (Fig. S8B), suggesting that the surface modification is crucial for specific binding to the T4P filament. As we did not detect filaments that were covered by the beads, we suggest that they only bind to a specific part of the filament, which is most likely the tip of the pilus.

**Fig. 6.**
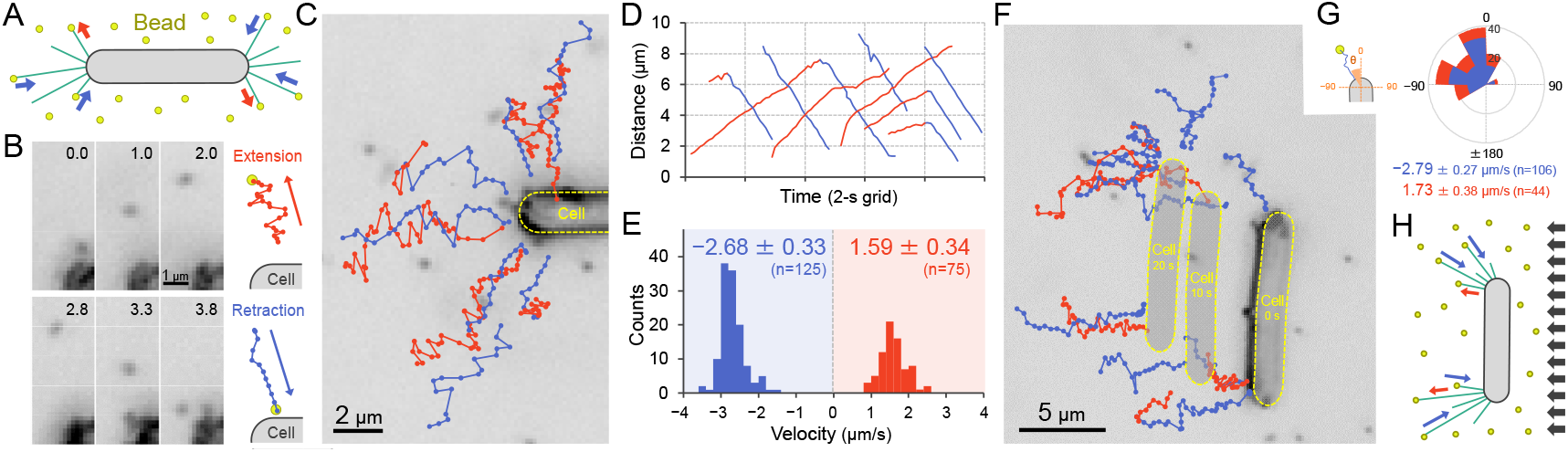
Visualization of T4P dynamics through nano-beads. (A) Schematic of the beads’ assay. Sulfate beads 200 nm in diameter were added to the cells on a glass surface. (B) Typical trajectories of beads. *Left*: a series of images. Time was presented at the right upper corner of the image. *Right*: trajectories of the bead moving away from and towards the cell. (C) Trajectories of beads. Red and blue indicate bead movement away from and towards the cell, respectively. The bead trajectories with a time interval of 0.1 s were overlaid onto the cell image. (D) Time course of the distance between the bead and a cell pole. The data came from Panel C. (E) Distribution of the bead velocity. The velocity was measured by the time course of bead displacement. The movement towards the cell was measured as a negative value. The average and SD of the plus and minus regions are presented (N = 200 in 12 cells). (F) Trajectories of beads during negative phototaxis. Lateral light illumination was applied from the right side of the image. The cell perpendicular to the lateral light axis is presented. The cell showed directional movement towards the left side of the image, and the cell position is presented by the dashed yellow lines every 10 s. The trajectories of beads with a time interval of 0.1 s were overlaid onto the cell image. (G) Distribution of the bead angle. *Left*: Schematic of the angle definition. The cells perpendicular to the lateral light axis were used for analysis. *Right*: The angle formed by the longer cell axis and the bead trajectories around the cell pole at the upper side were measured (N = 150 in 10 cells). (H) Schematic of T4P dynamics during negative phototaxis of the cell perpendicular to the lateral light axis. T4P was asymmetrically activated on the other side of the light source.

We also visualized the T4P dynamics through beads during negative phototaxis (Fig. 6F and Movie S15). Here, only cells that were aligned perpendicular to the optical axis of lateral light were used for data analyses. The beads were retracted only at the cell pole facing the opposite side of the light source, which is the leading pole in negative phototaxis (Fig. 6G). This suggests asymmetric activation of T4P at one of the two cell poles (Fig. 6H).

## Discussion

Here, we demonstrated that directional movement in phototaxis can be reversed by illumination with lateral light of another wavelength on a short time scale (Fig. 2 and Fig. 3). In bacterial twitching motility, cellular movement is driven by repeated cycles of extension and retraction of T4P (Fig. 4 and Fig. 6). In rod-shaped bacteria, such as *Myxococcus*, the T4P machinery is localized at cell poles and generates a directional bias presumably by the force generated at the leading pole along the longer axis of the cell (27–29). Here, we show that rod-shaped *T. vulcanus* cells showed directional movement even if the cell orientation was perpendicular to the lateral light axis (Fig. 5). This behaviour can be explained by asymmetric activation of T4P for the long cell axis at the pole on one side of the rod (Fig. 6). Such localization of pili was supported by our bead assay (Fig. 6F-H), which showed localized retraction of beads on the cell side that did not face the light source in negative phototaxis. Local activation of T4P along the shorter axis of the cell would require a specific signalling system with high spatial resolution compared to the well-known pole-to-pole regulation by *Myxococcus* and *Pseudomonas* (27–30). In these bacteria, a dynamic change in the localization of the motor protein PilB between both poles leads to a directional switch in motility (28, 30). The localization of PilB also seems to govern the direction of movement in *Synechocystis* phototaxis (31). Cell polarity for the short axis of the *T. vulcanus* cells might result in localization of the PilB protein to the leading side of the cell, driving directional movement even if the cell is oriented lengthwise with respect to the light source.

It remains an open question what the photoreceptor for positive phototaxis is. We showed that the gene disruption mutants of all photoreceptors identified to date for blue-to-green light still exhibited positive phototaxis (Fig. 3), suggesting that another unrecognized photoreceptor for positive phototaxis might remain to be identified in this species. In a wild isolate of *Synechococcus elongatus*, the cyanobacteriochrome PixJ was reported to be responsible for directional light sensing in phototaxis (9). A homologue of the PixJ protein was also shown to localize at cell poles in a closely related *Thermosynechococcus* species (32). However, the *pixJ* orthologue is deleted from the genome of our WT strain (33). Recent work reported that the GAF domain-containing photoreceptor PixJ is not responsible for the phototactic motility of the filament-forming cyanobacterium *Nostoc punctiforme* (34). The authors hypothesized that the localized proton motive force is the first input for directional light sensing (34). Considering the micro-optic effects in rod-shaped cyanobacteria (Fig. S10) (9), the spatial difference in light intensity between cell poles or different sides of the cell may be a universal key feature of T4P-dependent phototaxis.

How could c-di-GMP regulate the motility direction of *T. vulcanus* phototaxis? The 30–60 sec delay of the cellular response upon on/off illumination with blue light under background green light might result from the modulation of intracellular c-di-GMP levels or the activation of other downstream events. Considering the fast response of the directional switching of *T. vulcanus* motility in one minute, we assume that toggle regulation does not involve transcriptional regulation (35). As additional blue illumination has the same effect on phototaxis irrespective of the position of the blue light source (Fig. S6), we also assume that the increased global cellular pool of c-di-GMP under these conditions regulates the switch in movement.

In *Synechocystis*, multiple signalling networks seem to be integrated via PATAN-REC proteins, which interact with PilB1 (36, 37). We have not yet assessed the localization of the PilB protein, but we observed that the active pilus filaments face the direction of cell movement (Fig. 6B). Notably, the MshEN domain in cyanobacterial PilB proteins is a potential c-di-GMP binding domain despite the lack of experimental evidence (38). The switching between negative and positive phototaxis could be mediated by the binding and unbinding of c-di-GMP to PilB, respectively. We do not surmise that c-di-GMP binding would activate PilB-mediated pili extension, as claimed in other bacteria (39, 40), because we did not observe a change in pilus dynamics under green and green/blue light illumination (Fig. S9). Therefore, we hypothesize that c-di-GMP binding to PilB could modify the affinity of its binding to other interaction partner(s), leading to the different localization of PilB regarding the incident light vector. Blue light increases the intracellular c-di-GMP content and represses motility in *Synechocystis* on phototaxis plates (41), although short-term effects have not yet been explored. Whether c-di-GMP could induce a similar directional switching in other cyanobacterial species remains to be clarified. Notably, *Synechocystis* also switches from positive to negative phototaxis on a relatively short time scale by a different mechanism (36). High-intensity blue light is sensed by the BLUF-domain PixD photoreceptor. Upon excitation, the binding of PixD to the specific PATAN-domain CheY-like response regulator (PixE) is inhibited, and free PixE binds to PilB1, inducing a switch from red light-dependent positive to negative phototaxis (36). In *T. vulcanus*, the lack of pixD did not lead to directional switching (Fig. 3B). Thus, we suppose that different cyanobacteria have evolved specific photoreceptors and response mechanisms to control the switch between negative and positive phototaxis, presumably depending on their natural habitat and the ecophysiological context.

*T. vulcanus* was initially isolated from a mat in a hot spring in Japan (42). We previously reported that in multicellular cyanobacterial communities, the ratio between blue and green light changes from surface (blue rich) to green-rich within a culture due to pigment absorption. Thus, through the concerted action of the SesABC photoreceptor system, c-di-GMP production would be activated at the surface, whereas within a cyanobacterial community, low cellular c-di-GMP levels would predominate (22). The current study suggests that in a biofilm or a mat, cells move away from the blue light-rich surface area, whereas they move upwards under green-rich conditions within the community (Fig. S11). Such behaviour would lead to a dynamic circulation of *Thermosynechococcus* species inside a microbial mat under solar irradiance (43). The high light/short wavelength-induced downward migration and green light-induced upward migration of cyanobacteria in microbial mats are well documented (44–47). Future work on spatiotemporal processes in phototrophic mats can provide insights into the yet unknown molecular mechanisms of photomovements in ecophysiologically relevant niches.

## Materials and Methods

### Strains and culture conditions

The motile strains (WT) of *T. vulcanus* and its mutants were grown in BG-11 medium (48) containing 20 mM TES (pH 7.5) as a buffer in moderate light (20 μmol m^-2^ s^-1^) at 45 °C with shaking to an optical density of approximately 0.5–1.0 at 750 nm. The WT showing clear positive phototaxis on agar plates of BG11 was reisolated from the original strain NIES-2134 in the Microbial Culture Collection at the National Institute for Environmental Studies (NIES, https://mcc.nies.go.jp/) and used here as the standard strain.

### Strain construction

All primers, plasmids, and strains used in this study are listed in Tables S1, S2 and S3, respectively. The plasmid for the disruption of the *sesC* gene was already reported in (17). For the construction of the other mutant strains, an antibiotic resistance cassette was introduced to cause a partial deletion within each ORF. The PCR fragments of a vector backbone, an antibiotic resistance cassette, and the upstream and downstream sequences of 2–3 kbp were assembled using assembly cloning (AQUA cloning) (49). Transformations of *T. vulcanus* were performed according to (50). For preparation and cultivation of the mutant strains, antibiotics were added to the medium at the following concentrations: chloramphenicol, 5 μg ml^-1^; kanamycin 80 μg ml^-1^; and spectinomycin plus streptomycin, 10 μg ml^-1^ plus 5 μg ml^-1^. Complete segregation of the mutant alleles in the multiple copies of the chromosomal DNA was verified by colony PCR.

### Optical microscopy and data analyses

Cell behaviour on the glass surface was visualized under an inverted microscope (IX83; Olympus) equipped with 10× objective lenses (UPLFLN10×2PH, NA0.3; Olympus), a CMOS camera (Zyla 4.2; Andor, or DMK33U174; Imaging Source), and an optical table (ASD-1510T; JVI, Japan). The position of the cell was visualized by infrared light from a halogen lamp with a bandpass filter (FBH850/40; Thorlabs) at a fluence rate of 1 μmol m^-2^ s^-1^. The heat-absorbing filter was removed from the optical axis of the halogen lamp to obtain a higher wavelength of light. The signal from lateral light illumination was removed by a bandpass filter (FBH850/40; Thorlabs) in a filter block turret. The microscope stage was heated at 45 °C with a thermoplate (TP-110R-100; Tokai Hit, Japan). Projections of the images were captured as greyscale images with the camera under 1-s resolution and converted into a sequential TIF file without any compression. All data were analysed by ImageJ 1.48v (rsb.info.nih.gov/ij/) and its plugins, TrackMate, particle tracker and multitracker.

For the bead assay, the cells and microbeads were visualized under an inverted microscope (IX83; Olympus) equipped with 40× objective lenses (LUCPLFLN40×PH, NA0.6; Olympus), a CMOS camera (Zyla 4.2; Andor, or DMK33U174; Imaging Source), and an optical table (ASD-1510T; JVI, Japan). The position of the cell and microbeads were visualized by a collimated blue-light LED (M450LP1; Thorlabs) through a dark-field condenser (U-DCD, NA0.8-0.92; Olympus) at a fluence rate of 200 μmol m^-2^ s^-1^. The microscope stage was heated at 45 °C with a thermoplate (TP-110R-100; Tokai Hit). Projection of the image was captured as greyscale images with the camera under 0.1-s resolution and converted into a sequential TIF file without any compression.

### Phototaxis on glass with lateral light from LEDs

All procedures were performed at 45 °C on a microscope stage heated with a thermoplate (TP-110R-100; Tokai Hit, Japan). The cell culture was poured into a tunnel chamber assembled by taping a coverslip (7), and both ends of the chamber were sealed with nail polish to keep from drying the sample. The position of the cell was visualized by infrared light from a halogen lamp with a bandpass filter (FBH850/40; Thorlabs) at a fluence rate of 1 μmol m^-2^ s^-1^. The cells were subjected to lateral light stimulus by an LED from the right side of the microscope stage at an angle of 5 degrees. White LEDs at 20 and 500 μmol m^-2^ s^-1^ were used as moderate and strong light stimuli for phototaxis, respectively. Blue, teal, green, orange, red, and far-red light were applied by a monochromatic LED, M450LP1, M490L4, M530L3, M625L3, and M730L4 (Thorlabs), respectively. The LED light was collimated by the condenser lens and combined by dichroic mirrors (FF470-Di01, FF509-FDi01, FF560-FDi01, FF685-Di02; Semrock) to apply multicoloured light simultaneously. The wavelength of the resultant light was measured by a spectrometer (BIM-6002A, BroLight, China). Light intensity was measured with a power metre (Q82017A; Advantest, Japan).

### Measurement of intracellular c-di-GMP concentration

Liquid cultures (1.5 ml) of WT and Δ*sesA* strain of *T. vulcanus* were transferred to three prewarmed 2 ml tubes. Two tubes were used for the total nucleotide extraction, and the third tube was used for the determination of the protein concentration. The tubes were incubated under 70 μmol m^-2^ s^-1^ blue, green, red (λ_max_=451, 528, 625 nm), or white LED light (Philips LED tube 16W830) illumination at 45 °C for 30 min. The cells were pelleted at 10,000 *g* at 4 °C for 30 sec. For nucleotide extraction, cell pellets were immediately resuspended in 200 μl extraction solution (acetonitrile/methanol/water 2:2:1 [v/v/v]), vortexed for 5 sec, and heated at 95 °C for 10 min. Samples were snap-cooled on ice and incubated for 15 min. The samples were centrifuged at 21,000 *g* at 4 °C for 10 min, and the supernatant was collected. The above extraction was repeated twice with 200 μl extraction solution each, without heat treatment. The combined supernatants (~600 μl) were incubated at −20 °C overnight to precipitate proteins in the samples. The samples were centrifuged at 21,000 *g* at 4 °C for 10 min, and the supernatant was vacuum dried using SpeedVac at 42 °C. Quantification of c-di-GMP was performed by HPLC/MS/MS analysis, as previously described (51). For quantification of the total protein content, the pelleted cells were stored at −20 °C. The de-frozen cells were suspended in 50 μl phosphate-buffered saline. A glass bead mix (0.1–0.11 and 0.25–0.5 mm) of ~0.7 volume was added. The tubes were vortexed vigorously for 60 sec, snap-cooled at −80 °C, and heated to 40 °C for 10 min. The disruption was repeated once and incubated at RT until the beads were sedimented. Two microlitres of the supernatant was used for protein quantification using the Direct Detect system (Merck Millipore). The acquired concentrations of intracellular c-di-GMP were normalized to the total protein content.

### Bead’s assay for visualizing T4P dynamics

Fluorescent polystyrene beads 0.2 μm in size (FluoSpheres sulfate microspheres F8848, carboxylate-modified microspheres F8811, amine-modified microsphere F8764; Thermo Fisher) were diluted 300 times to 0.02% (wt/vol) in BG11 and used for the bead assay, as previously described (7). A coverslip was coated with 0.2% (vol/vol) collodion in isoamyl acetate and air-dried before use. The cell culture was poured into a tunnel chamber assembled by taping a coverslip. After incubation at 45 °C for 2 min on the microscope stage, the cells were subjected to vertical illumination from blue-light LED through a dark-field condenser at a fluence rate of 200 μmol m^-2^ s^-1^. After illumination for 2 min, fluorescent beads were added to the sample chamber, and their movement was visualized by blue light illumination at 0.1-s intervals. Lateral illumination from green-light LEDs was applied for 2 min before adding the beads if needed.

### Electron Microscopy

Samples bound to the grids were stained with 2% (wt/vol) ammonium molybdate and observed by transmission electron microscopy, as previously described (7). Carbon-coated EM grids were prepared by a vacuum evaporator (VE-2012; Vacuum Device, Japan). Cells were placed on an EM grid and kept at 45 °C in moderate light (20 μmol m^-2^ s^-1^) for 3 min. The cells were chemically fixed with 1% (vol/vol) glutaraldehyde in BG-11 for 10 min at RT. After washing three times with BG-11, the cells were stained with 2% ammonium molybdate and air-dried. Samples were observed under a transmission electron microscope (JEM-1400, JEOL) at 100 kV. The EM images were captured by a charge-coupled device (CCD) camera.

## Acknowledgements

The authors thank Dr. Masahiko Ikeuchi (The University of Tokyo) for supplying the wild-type strain and Dr. Heike Bähre (Medical School Hannover) for c-di-GMP measurements. This study was supported in part by KAKENHI (16H06230, 20H05543, 21K07020) to DN, by funds from the Nakajima Foundation, the Noguchi Institute to DN, by KAKENHI (17K15244) to GE, by the German Science Foundation (WI2014/7-1, WI2014/8-1 and in frame of the SFB1381 - 403222702-SFB 1381 (A2)) to AW. GE is supported by the EMBO postdoctoral fellowship (ALTF 274-2017) and JSPS Overseas Research Fellowships.

## Author contributions

All authors designed the research, DN and GE performed the experiments, DN and GE performed the data analyses, and all authors wrote the paper.

## Data and materials availability

All data needed to evaluate the conclusions in the paper are present in the paper or the supplementary materials.

