## Supplementary Information for "*Thermosynechococcus* switches the direction of phototaxis by a c-di-GMP dependent process with high spatial resolution"

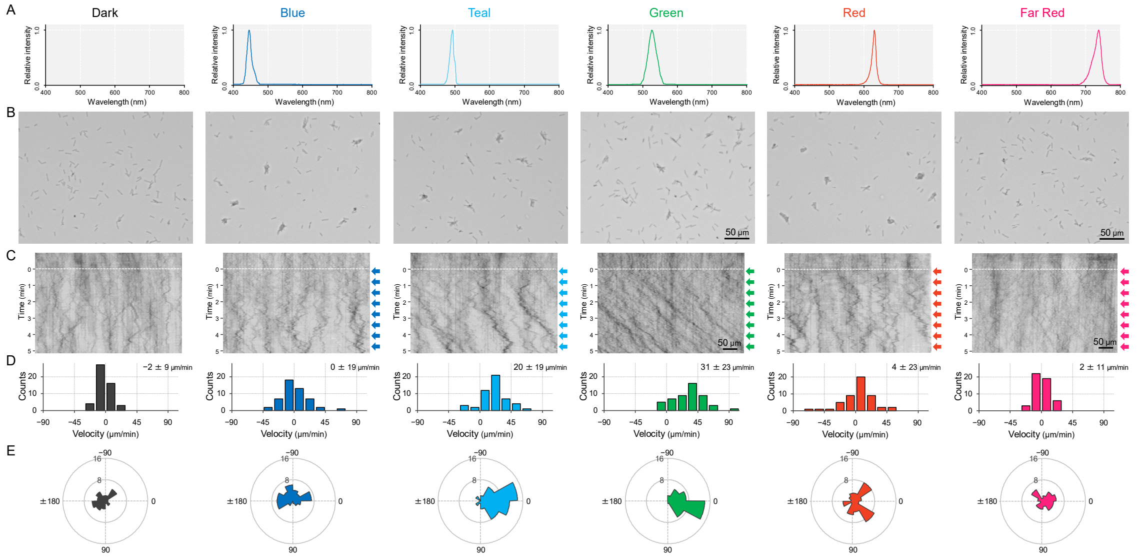


**Fig. S1.** Cell movement after applying a monochromatic light source. (A) Spectra of each LED used for stimulation. No lateral illumination in the dark condition. Infrared light was used for cell observation from a halogen lamp through a bandpass filter (see more detail in Fig. 2AB and Materials and Methods). (B) Cell images at 4 min after the lateral light was turned on. Lateral light from each LED was adjusted at a fluence rate of 70 μmol $m^{-2}$ $s^{-1}$ and applied from the right side of the image. (C) Kymograph of cell movements along the optical axis of lateral illumination. Directional movements of cells are presented by the tilted lines over time. Lateral light illumination was turned on at time 0, presented as a dashed white line. (D) Histograms of the cell velocity along the lateral light axis. Cell movements towards the light source are shown as a positive value. (E) Rose plots. The moving direction of a cell that translocated more than 6 μm $min^{-1}$ was counted. Angle 0 was the direction towards the lateral light source. The cell displacement for a duration of 1 minute was measured at 4 min after lateral illumination was turned on (N = 50 cells).

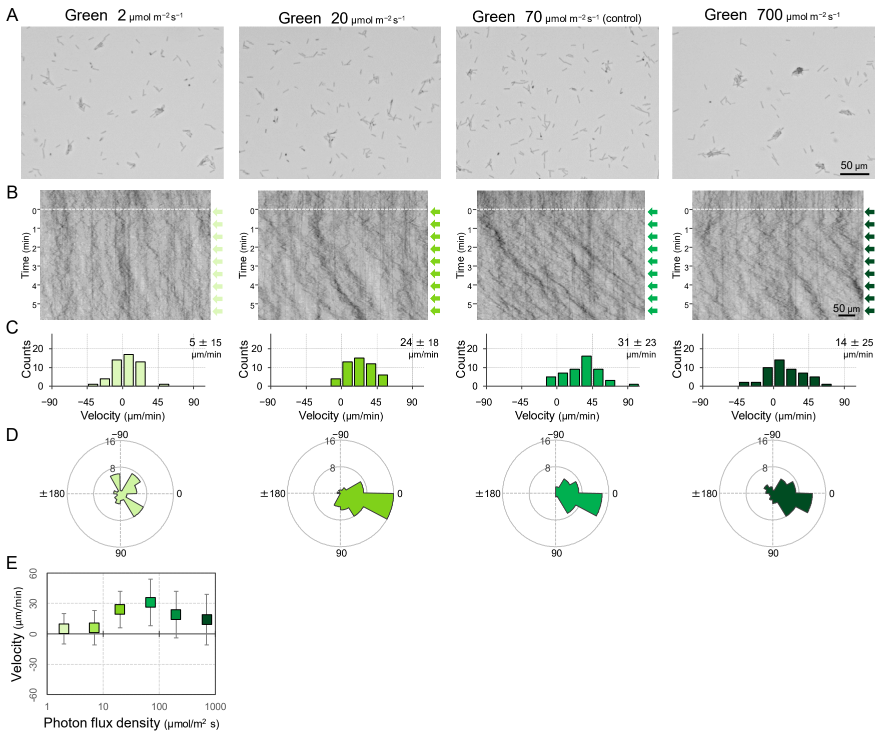


**Fig. S2.** Cell movement at various fluence rates of a lateral green light. (A) Cell images at 4 min after the lateral light was turned on. (B) Kymograph of cell movements along the optical axis of lateral illumination. Directional movements of cells are presented by the tilted lines over time. Lateral light illumination was turned on at time 0, presented as a dashed white line. (C) Histograms of the cell velocity along the lateral light axis. Cell movements towards the light source are shown as a positive value. (D) Rose plots. The moving direction of a cell that translocated more than 6 μm $min^{-1}$ was counted. Angle 0 was the direction towards the lateral light source. The cell displacement for a duration of 1 minute was measured at 4 min after lateral illumination was turned on (N = 50 cells). (E) Effects of the fluence rate on the phototactic behaviour of cells. The average and standard deviation (SD) of the cell velocity along the light axis are presented (N = 50).

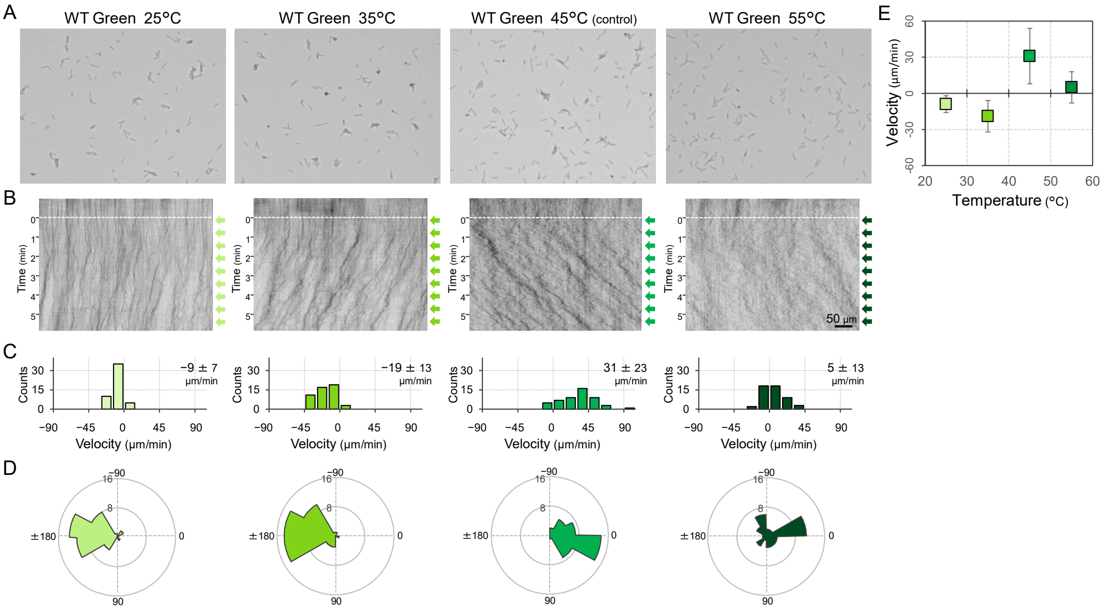


**Fig. S3.** Phototactic behaviour of WT cells at various temperatures. Lateral light from a single LED was adjusted to a fluence rate of 70 µmol $m^{-2}$ $s^{-1}$ and applied from the right side of the image. (A) Cell image at 4 min after the lateral light was turned on. (B) Kymograph of cell movements along the optical axis of lateral illumination. Directional movements of cells are presented by the tilted lines over time. Lateral light illumination was turned on at time 0, presented as a dashed white line. (C) Histograms of the cell velocity along the lateral light axis. Cell movements towards the light source are shown as a positive value. (D) Rose plots. The moving direction of a cell that translocated more than 6 µm $min^{-1}$ was counted. Angle 0 was the direction towards the lateral light source. The cell displacement for a duration of 1 minute was measured at 4 min after lateral illumination was turned on (N = 50 cells). (E) Effects of the fluence rate on the phototactic behaviour of WT cells. The average and standard deviation (SD) of the cell velocity along the light axis are presented (N = 50).

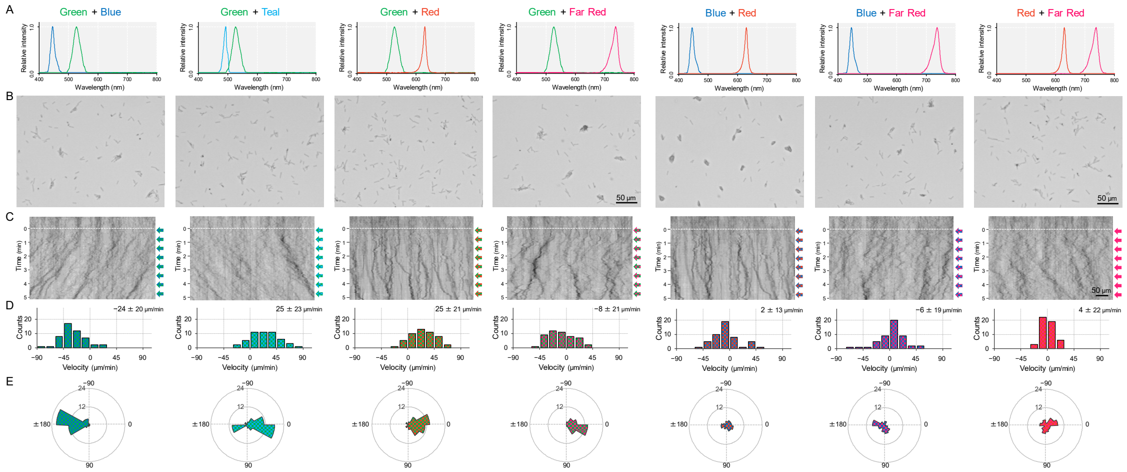


**Fig. S4.** Cell movement after applying a dichromatic light source. (A) Spectra of two LEDs used for stimulation. (B) Cell images at 4 min after the lateral light was turned on. Lateral light from a single LED was adjusted at a fluence rate of 70 μmol $m^{-2}$ $s^{-1}$ and applied from the right side of the image. (C) Kymograph of cell movements along the optical axis of lateral illumination. Directional movements of cells are presented by the tilted lines over time. Lateral light illumination was turned on at time 0, presented as a dashed white line. (D) Histograms of the cell velocity along the lateral light axis. Cell movements towards the light source are shown as a positive value. (E) Rose plots. The moving direction of a cell that translocated more than 6 μm $min^{-1}$ was counted. Angle 0 was the direction towards the lateral light source. The cell displacement for a duration of 1 minute was measured at 4 min after lateral illumination was turned on (N = 50 cells).

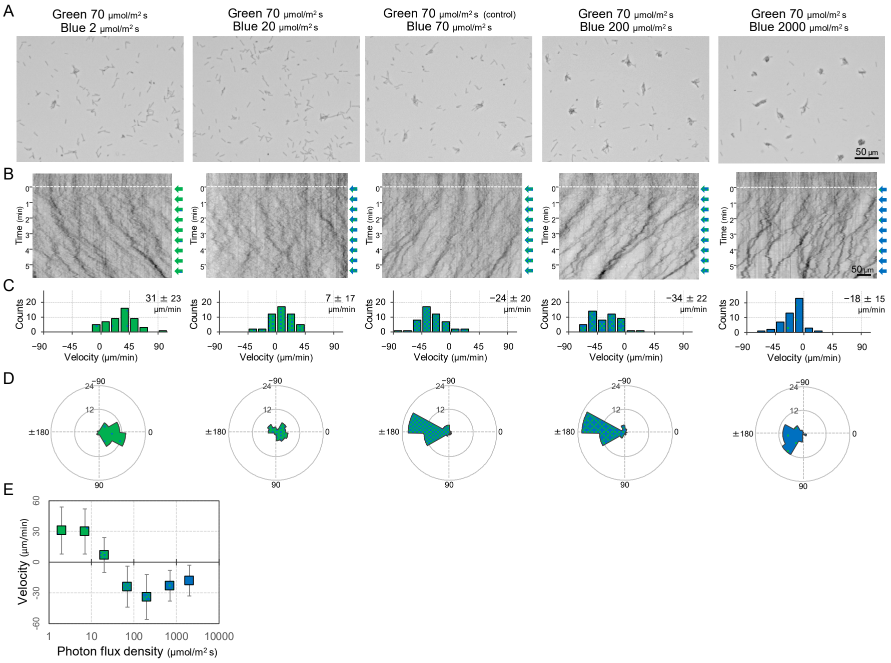


**Fig. S5.** Dose dependency of blue light to induce negative phototaxis. Lateral blue light was adjusted at various fluence rates, while lateral green light was fixed at 70 μmol $m^{-2}$ $s^{-1}$, as presented at the top. (A) Cell images at 4 min after the lateral light was turned on. (B) Kymograph of cell movements along the optical axis of lateral illumination. Directional movements of cells are presented by the tilted lines over time. Lateral light illumination was turned on at time 0, presented as a dashed white line. (C) Histograms of the cell velocity along the lateral light axis. Cell movements towards the light source are shown as a positive value. (D) Rose plots. The moving direction of a cell that translocated more than 6 μm $min^{-1}$ was counted. Angle 0 was the direction towards the lateral light source. The cell displacement for a duration of 1 minute was measured at 4 min after lateral illumination was turned on (N = 50 cells). (E) Effects of the fluence rate of blue light to induce negative phototaxis. The average and standard deviation (SD) of the cell velocity along the light axis are presented (N = 50).

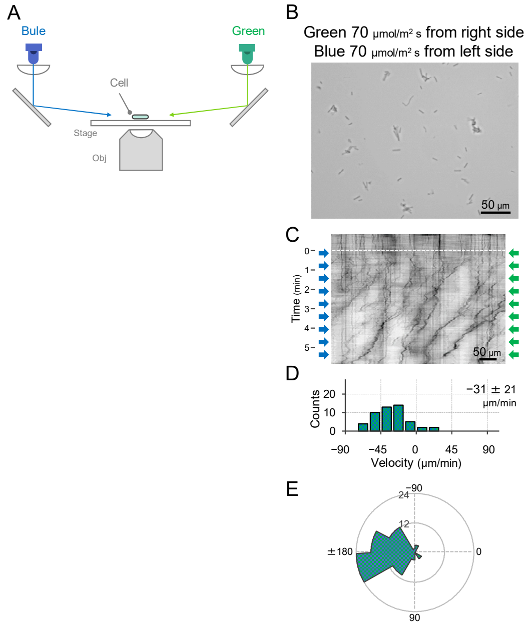


**Fig. S6.** Effect of blue light illumination from the other side of green light on phototaxis. (A) Schematic of the lateral light illumination. Blue and green light were applied from the left and right sides of the image at 70 μmol $m^{-2}$ $s^{-1}$, respectively. (B) Cell images at 4 min after the lateral light was turned on. (C) Kymograph of cell movements along the optical axis of lateral illumination. Directional movements of cells are presented by the tilted lines over time. Lateral light illumination was turned on at time 0, presented as a dashed white line. (D) Histograms of the cell velocity along the lateral light axis. Cell movements towards the light source are shown as a positive value. The average and standard deviation (SD) of the cell velocity along the light axis are presented (N = 50). (E) Rose plots. The moving direction of a cell that translocated more than 6 μm $min^{-1}$ was counted. Angle 0 was the direction towards the lateral light source of the green light. The cell displacement for a duration of 1 minute was measured at 4 min after lateral illumination was turned on (N = 50 cells).

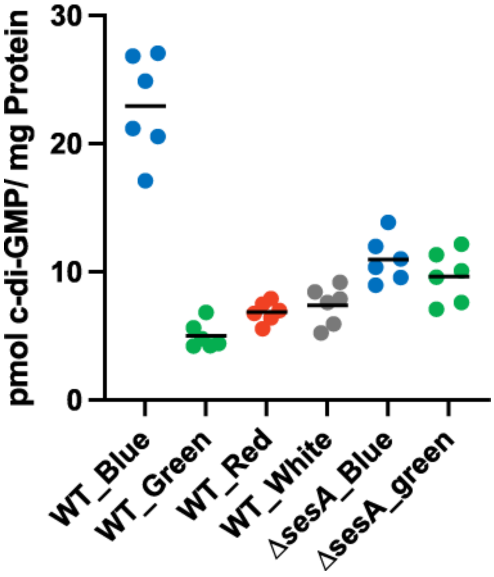


**Fig. S7.** Intracellular c-di-GMP concentrations in WT and Δ*sesA*. The cells were cultivated under blue, green, red, or white light illumination for 30 min, and c-di-GMP was extracted and quantified. The shown data are biological triplicates with technical duplicates, and the mean values are given with the bars.

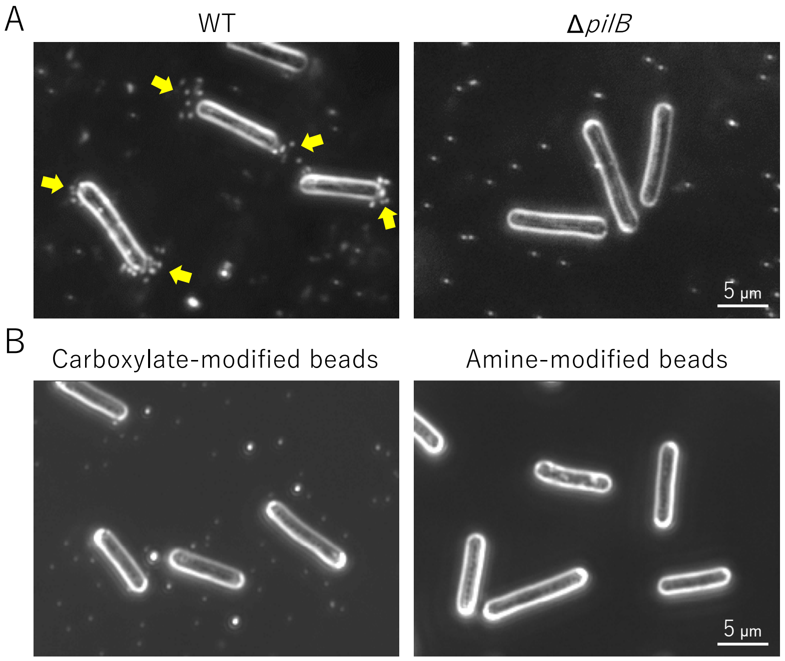


**Fig. S8.** Optimization of beads assay. (A) T4P-dependent bead accumulation at the cell pole. Cell images were observed by dark-field microscopy 2 min after adding 200 nm diameter sulfate beads. Yellow indicates bead accumulation at the cell pole. (B) Preference of bead accumulation. Dark-field images 2 min after adding 200 nm diameter carboxylate- and amine-modified beads are presented.

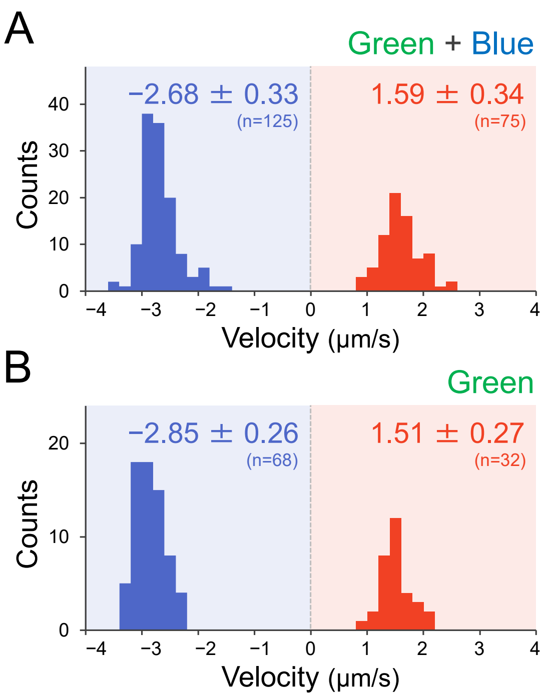


**Fig. S9.** Effect of the illumination of green and green blue light on T4P dynamics. T4P dynamics were visualized through 200 nm diameter sulfate beads . The velocity of bead movement was measured by the time course of bead displacement. The movement towards the cell was measured as a negative value. The average and SD of the plus and minus regions are presented. (A) Green light at 70 μmol $m^{-2}$ $s^{-1}$ (N = 200 in 12 cells). The same data are presented in Fig. 6E as a control. (B) Green and blue light at each 70 μmol $m^{-2}$ $s^{-1}$ (N = 100 in 10 cells).

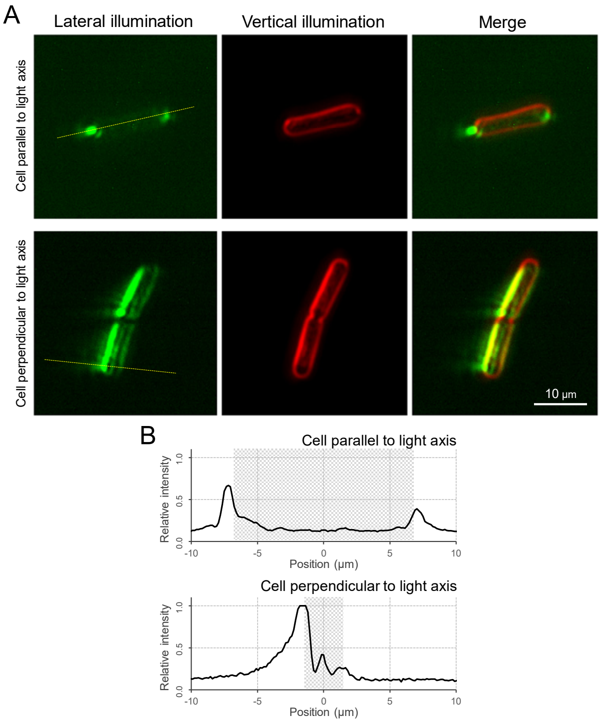


**Fig. S10.** Micro-optics effect. (A) Cell image via optical microscopy. *Left*: Cell image visualized by the lateral illumination of green light from the right side of the image. *Middle*: Cell outline visualized by the vertical illumination of green light. *Right*: Merged image. The cells attached on the glass surface parallel and perpendicular to the lateral light axis are presented at the *top* and *bottom*, respectively. (B) Intensity profiles. Light intensity along the yellow dashed line in Panel A is presented. The cell positions were marked by the grey regions.

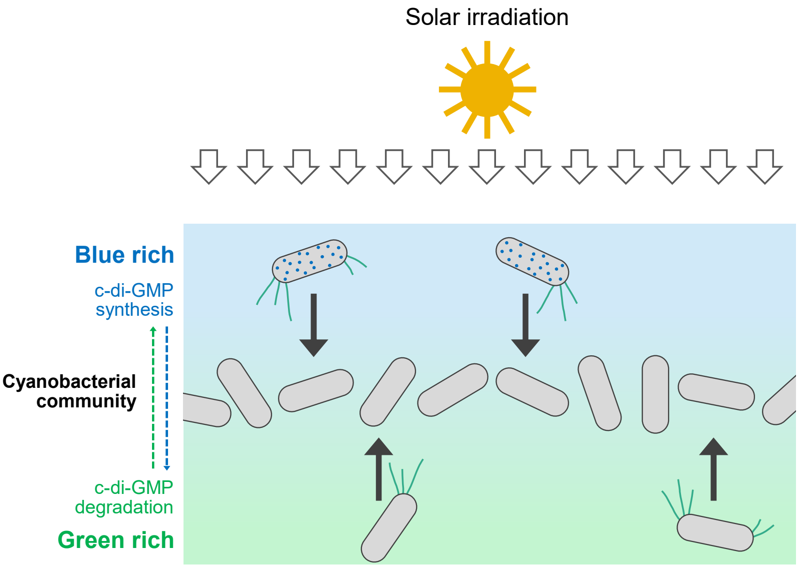


**Fig. S11.** Schematic illustration of cellular c-di-GMP levels and cell migration within a dense cyanobacterial community under solar irradiation. Blue light is rich at the surface of such a community (e.g., a phototrophic mat in a hot spring). Synthesis of c-di-GMP induces downward migration at the surface, while degradation of c-di-GMP induces upward migration inside.

**Table S1**: Oligonucleotides used in this study

| Name | Sequence (5'-3') | Purpose |
|---|---|---|
| pUC19-5R_tvsesA | ATGATACGCGGGTGATTACCGAGCTCGAATTCAC | Δ*sesA* |
| pUC19-6F_tvsesA | GTGGCAAAACCACCTTTGCAAGCTTGGCGTAATC | |
| tvsesA-1F_pUC19 | GAATTCGAGCTCGGTAATCACCCGCGTATCATTG | |
| tvsesA-2R_Cm | TTCTATCAGCTGTCCCATCAATCCCCGAAACTGC | |
| Cm-3F_tvsesA | AGTTTCGGGGATTGATGGGACAGCTGATAGAAAC | |
| Cm-4R_tvsesA | ATGGGTGATATTGGCTTTTATCAGGCTCTGGGAG | |
| tvsesA-3F_Cm | CCCAGAGCCTGATAAAAGCCAATATCACCCATGC | |
| tvsesA-4R_pUC19 | TTACGCCAAGCTTGCAAAGGTGGTTTTGCCACAG | |
| pUC19-7R_tvsesB | TCAGGGGAGTCCAAAATACCGAGCTCGAATTCAC | Δ*sesB* |
| pUC19-8F_tvsesB | ACGGATTGCCATTGGTTGCAAGCTTGGCGTAATC | |
| tvsesB-1F_pUC19 | GAATTCGAGCTCGGTATTTTGGACTCCCCTGATG | |
| tvsesB-2R_Km | AGCAGGGGAATTGTTACACAGACTCTTCTGTGAC | |
| Km-5F_tvsesB | CACAGAAGAGTCTGTGTAACAATTCCCCTGCTCG | |
| Km-6R_tvsesB | GTTCAAGGTCCTCTTGCCAACCTTTCATAGAAGGC | |
| tvsesB-3F_Km | TTCTATGAAAGGTTGGCAAGAGGACCTTGAACTG | |
| tvsesB-4R_pUC19 | TTACGCCAAGCTTGCAACCAATGGCAATCCGTAG | |
| pUC19-21R_tvcikA | AGATGCGATTGTCCATTACCGAGCTCGAATTCAC | Δ*cikA* |
| pUC19-22F_tvcikA | TTCATTGGGCTTACAGTGCAAGCTTGGCGTAATC | |
| tvcikA-1F_pUC19 | GAATTCGAGCTCGGTAATGGACAATCGCATCTGC | |
| tvcikA-2R_Cm | TTCTATCAGCTGTCCCGCTGATAGCTACTGCTAG | |
| Cm-18F_tvcikA | AGCAGTAGCTATCAGCGGGACAGCTGATAGAAAC | |
| Cm-19R_tvcikA | AGATATTCCCGCTGTTTTTATCAGGCTCTGGGAG | |
| tvcikA-3F_Cm | CCCAGAGCCTGATAAAAACAGCGGGAATATCTGG | |
| tvcikA-4R_pUC19 | TTACGCCAAGCTTGCACTGTAAGCCCAATGAACC | |
| pUC19-19R_tvpixD | GTTACCATTCGCAAGTTACCGAGCTCGAATTCAC | Δ*pixD* |
| pUC19-20F_tvpixD | GCTAGTCCTGAACCAATGCAAGCTTGGCGTAATC | |
| tvpixD-1F_pUC19 | GAATTCGAGCTCGGTAACTTGCGAATGGTAACCG | |
| tvpixD-2R_Cm | TTCTATCAGCTGTCCCAATCAGGCGATGTAGTCC | |
| Cm-15F_tvpixD | ACTACATCGCCTGATTGGGACAGCTGATAGAAAC | |
| Cm-16R_tvpixD | CAGCATTCCCGTAATGTTTATCAGGCTCTGGGAG | |
| tvpixD-3F_Cm | CCCAGAGCCTGATAAACATTACGGGAATGCTGTG | |
| tvpixD-4R_pUC19 | TTACGCCAAGCTTGCATTGGTTCAGGACTAGCTG | |
| pUC19-23R_tvtll1282 | TTTGGCATCTTCAAGGTACCGAGCTCGAATTCAC | Δ*LOV* |
| pUC19-24F_tvtll1282 | TGAGCAAAGGGATCATTGCAAGCTTGGCGTAATC | |
| tvtll1282-1F_pUC19 | GAATTCGAGCTCGGTACCTTGAAGATGCCAAAGC | |
| tvtll1282-2R_Cm | TTCTATCAGCTGTCCCAGCATAGCGTTCCATCTC | |

| | | |
|---|---|---|
| Cm-20F_tvtll1282 | GATGGAACGCTATGCTGGGACAGCTGATAGAAAC | |
| Cm-21R_tvtll1282 | AATAGGCTCAGGACATTTTATCAGGCTCTGGGAG | |
| tvtll1282-3F_Cm | CCCAGAGCCTGATAAAATGTCCTGAGCCTATTGG | |
| tvtll1282-4R_pUC19 | TTACGCCAAGCTTGCAATGATCCCTTTGCTCAGC | |
| pUC19-25R_tvHCP | TGAACTTTGCCCTTGTTACCGAGCTCGAATTCAC | *ΔOCP* |
| pUC19-26F_tvCTDH | GATAGGGAGATTGCATTGCAAGCTTGGCGTAATC | |
| tvHCP-1F_pUC19 | GAATTCGAGCTCGGTAACAAGGGCAAAGTTCAGC | |
| tvHCP-2R_Cm | TTCTATCAGCTGTCCCGCTCATCAAAACTGAGGG | |
| Cm-22F_tvHCP | CTCAGTTTTGATGAGCGGGACAGCTGATAGAAAC | |
| Cm-23R_tvCTDH | TCTTCAAAGGGTGGATTTTATCAGGCTCTGGGAG | |
| tvCTDH-1F_Cm | CCCAGAGCCTGATAAAATCCACCCTTTGAAGAGC | |
| tvCTDH-2R_pUC19 | TTACGCCAAGCTTGCAATGCAATCTCCCTATCCG | |
| pUC19-15R_tvpilB | CAGTTGAATCTGGGTTTACCGAGCTCGAATTCAC | *ΔpilB* |
| pUC19-16F_tvpilB | ACAGGTTTTGGAGGTCTGCAAGCTTGGCGTAATC | |
| tvpilB-1F_pUC19 | GAATTCGAGCTCGGTAAACCCAGATTCAACTGGG | |
| tvpilB-2R_Cm | TTCTATCAGCTGTCCCTTGTGGTACTGGCGTAAC | |
| Cm-11F_tvpilB | TACGCCAGTACCACAAGGGACAGCTGATAGAAAC | |
| Cm-12R_tvpilB | AGATCTTGAAGCGGGATTTATCAGGCTCTGGGAG | |
| tvpilB-3F_Cm | CCCAGAGCCTGATAAATCCCGCTTCAAGATCTTG | |
| tvpilB-4R_pUC19 | TTACGCCAAGCTTGCAGACCTCCAAAACCTGTAC | |
| pUC19-17R_tvpilT1 | TTTTTGCCAGCACCTTTACCGAGCTCGAATTCAC | *ΔpilT1* |
| pUC19-18F_tvpilT1 | TCTTGGCTGGCTTGATTGCAAGCTTGGCGTAATC | |
| tvpilT1-1F_pUC19 | GAATTCGAGCTCGGTAAAGGTGCTGGCAAAAAGC | |
| tvpilT1-2R_Cm | TTCTATCAGCTGTCCCAAGTCTGAACCACCGTTG | |
| Cm-13F_tvpilT1 | ACGGTGGTTCAGACTTGGGACAGCTGATAGAAAC | |
| Cm-14R_tvpilT1 | TGGCAAGCTGAATCGTTTTATCAGGCTCTGGGAG | |
| tvpilT1-3F_Cm | CCCAGAGCCTGATAAAACGATTCAGCTTGCCATC | |
| tvpilT1-4R_pUC19 | TTACGCCAAGCTTGCAATCAAGCCAGCCAAGAAG | |
| pUC19-11R_tvpilA1 | CTATTCTCTTTGCAGGTACCGAGCTCGAATTCAC | *ΔpilA1* |
| pUC19-12F_tvpilA1 | TGGTTTGGTGCCAGTATGCAAGCTTGGCGTAATC | |
| tvpilA1-1F_pUC19 | GAATTCGAGCTCGGTACCTGCAAAGAGAATAGCG | |
| tvpilA1-7R_Cm | TTCTATCAGCTGTCCCAGTTGCCGTCTGCTACAG | |
| Cm-32F_tvpilA1 | GTAGCAGACGGCAACTGGGACAGCTGATAGAAAC | |
| Cm-6R_tvpilA1 | GCAGTTGCACTATTGCTTTATCAGGCTCTGGGAG | |
| tvpilA1-3F_Cm | CCCAGAGCCTGATAAAGCAATAGTGCAACTGCTC | |
| tvpilA1-4R_pUC19 | TTACGCCAAGCTTGCATACTGGCACCAAACCATG | |
| pUC19-13R_tvHfq | CAGCGGGATGTGAATATACCGAGCTCGAATTCAC | *Δhfq* |
| pUC19-14F_tvHfq | AAATGCGGCTTTCCTATGCAAGCTTGGCGTAATC | |

| | |
|---|---|
| tvHfq-1F_pUC19 | GAATTCGAGCTCGGTATATTCACATCCCGCTGTG |
| tvHfq-2R_Cm | TTCTATCAGCTGTCCCTACTTGGCGAATGCTAGG |
| Cm-9F_tvHfq | TAGCATTCGCCAAGTAGGGACAGCTGATAGAAAC |
| Cm-10R_tvHfq | ACCTCGGTGCTATCTTTTTATCAGGCTCTGGGAG |
| tvHfq-3F_Cm | CCCAGAGCCTGATAAAAAGATAGCACCGAGGTTG |
| tvHfq-4R_pUC19 | TTACGCCAAGCTTGCATAGGAAAGCCGCATTTGC |

**Table S2**: Plasmids used in this study

| Name | Description | Reference/Source |
|---|---|---|
| pUC19-ΔtvsesA_Cm | pUC19-based construct for knock-out of *sesA* gene (NIES2134_109940) containing a chloramphenicol resistance cassette flanked by ~2500 bp regions upstream and downstream of *sesA,* resulting in ~ 2000 bp deletion within ORF; Cm$^R$, Amp$^R$ | This study |
| pUC19-ΔtvsesB_Km | pUC19-based construct for knock-out of *sesB* gene (NIES2134_119260) containing a kanamycin resistance cassette flanked by ~2500 bp regions upstream and downstream of *sesB,* resulting in ~ 2000 bp deletion within ORF; Km$^R$, Amp$^R$ | This study |
| pS-Δtlr0911_Sp(F) | pCR-Script-based construct for knock-out of *sesC* gene (NIES2134_110090) containing a spectinomycin/streptomycin resistance cassette flanked by ~1500 bp regions upstream and downstream of *sesC,* resulting in ~ 3400 bp deletion within ORF; Sp$^R$/Sm$^R$, Amp$^R$ | (14) |
| pUC19-ΔtvcikA_Cm | pUC19-based construct for knock-out of *cikA* gene (NIES2134_110210) containing a chloramphenicol resistance cassette flanked by ~2500 bp regions upstream and downstream of *cikA,* resulting in ~ 500 bp deletion within ORF; Cm$^R$, Amp$^R$ | This study |
| pUC19-ΔtvpixD_Cm | pUC19-based construct for knock-out of *pixD* gene (NIES2134_124540) containing a chloramphenicol resistance cassette flanked by ~2500 bp regions upstream and downstream of *pixD,* resulting in ~ 100 bp deletion within ORF; Cm$^R$, Amp$^R$ | This study |
| pUC19-Δtvtll1282_Cm | pUC19-based construct for knock-out of *LOV* gene (*tll1282;* NIES2134_112750) containing a chloramphenicol resistance cassette flanked by ~2500 bp regions upstream and downstream of *cikA,* resulting in ~ 2000 bp deletion within ORF; Cm$^R$, Amp$^R$ | This study |
| pUC19-ΔtvHCPCTDH_Cm | pUC19-based construct for knock-out of *OCP* genes (HCP; NIES2134_112880 and CTDH; | This study |

| | | |
|---|---|---|
| | NIES2134_112890) containing a chloramphenicol resistance cassette flanked by ~2500 bp regions upstream and downstream of *OCP,* resulting in the deletion of the latter half of *HCP* ORF, the intergenic region, and the first half of *CTDH* ORF; $Cm^R$, $Amp^R$ | |
| pUC19-$\Delta$tvpilB_Cm | pUC19-based construct for knock-out of *pilB* gene (NIES2134_124120) containing a chloramphenicol resistance cassette flanked by ~2500 bp regions upstream and downstream of *pilB,* resulting in ~ 600 bp deletion within ORF; $Cm^R$, $Amp^R$ | This study |
| pUC19-$\Delta$tvpilT1_Cm | pUC19-based construct for knock-out of *pilT1* gene (NIES2134_124130) containing a chloramphenicol resistance cassette flanked by ~2500 bp regions upstream and downstream of *pilT1,* resulting in ~ 600 bp deletion within ORF; $Cm^R$, $Amp^R$ | This study |
| pUC19-$\Delta$tvpilA1_Cm | pUC19-based construct for knock-out of *pilA1* gene (NIES2134_108920) containing a chloramphenicol resistance cassette flanked by ~2500 bp regions upstream and downstream of *pilA1,* resulting in ~ 550 bp deletion within ORF and the promoter region; $Cm^R$, $Amp^R$ | This study |
| pUC19-$\Delta$tvHfq_Cm | pUC19-based construct for knock-out of *hfq* gene (NIES2134_124130) containing a chloramphenicol resistance cassette flanked by ~2500 bp regions upstream and downstream of *hfq,* resulting in ~ 10 bp deletion within ORF; $Cm^R$, $Amp^R$ | This study |

**Table S3**: Strains used in this study

| Name | Description | Reference/Source |
|---|---|---|
| *Thermosynechococcus vulcanus* WT_P | Isolated as wild type showing positive phototaxis toward moderate white light from NIES-2134 | Whole genome: AP018202 |
| Δ*sesA* | *sesA* knockout mutant ($Cm^R$) | This study |
| Δ*sesB* | *sesB* knockout mutant ($Km^R$) | This study |
| Δ*sesC* | *sesC* knockout mutant ($Sp^R$/$Sm^R$) | (17) |
| Δ*cikA* | *cikA* knockout mutant ($Cm^R$) | This study |
| Δ*pixD* | *pixD* knockout mutant ($Cm^R$) | This study |
| Δ*LOV* | *LOV* knockout mutant ($Cm^R$) | This study |
| Δ*OCP* | *OCP* knockout mutant ($Cm^R$) | This study |
| Δ*pilB* | *pilB* knockout mutant ($Cm^R$) | This study |
| Δ*pilT1* | *pilT1*knockout mutant ($Cm^R$) | This study |
| Δ*pilA1* | *pilA1*knockout mutant ($Cm^R$) | This study |
| Δ*hfq* | *hfq* knockout mutant ($Cm^R$) | This study |

### Captions for Supplementary Movies

**Movie S1. Positive phototaxis.** Lateral illumination of green light was turned on at time 0 from the right side of the movie. WT cell at 45 °C. Area 750 × 563 μm.

**Movie S2. Microcolony formation.** Lateral illumination of blue light was turned on at time 0 from the right side of the movie. WT cell at 45 °C. Area 750 × 563 μm.

**Movie S3. Phototaxis at low temperature.** Lateral illumination of green light was turned on at time 0 from the right side of the movie. WT cell at 25 °C. Area 750 × 563 μm.

**Movie S4. Negative phototaxis.** Lateral illumination of green and blue light was turned on at time 0 from the right side of the movie. WT cell at 45 °C. Area 750 × 563 μm.

**Movie S5. On-off control of positive phototaxis.** Lateral illumination of green light was applied with 4 min intervals from the right side of the movie. WT cell at 45 °C. Area 750 × 563 μm.

**Movie S6. On-off control of negative phototaxis.** Lateral illumination of green and blue light was applied with 4 min intervals from the right side of the movie. WT cell at 45 °C. Area 750 × 563 μm.

**Movie S7. Directional switch of phototaxis in WT.** Lateral illumination was turned on at time 0 from the right side of the movie. The illumination was applied with three phases in the order of green, green/blue, and green with a time interval of 4 min. WT cell at 45 °C. Area 750 × 563 μm.

**Movie S8. Cell behaviour in Δ*sesA* mutant.** Lateral illumination was turned on at time 0 from the right side of the movie. The illumination was applied with three phases in the order of green, green/blue, and green with a time interval of 4 min. Δ*sesA* cell. Area 750 × 563 μm.

**Movie S9. Cell behaviour in Δ*sesC* mutant.** Lateral illumination was turned on at time 0 from the right side of the movie. The illumination was applied with three phases in the order of green, green/blue, and green with a time interval of 4 min. Δ*sesC* cell. Area 750 × 563 μm.

**Movie S10. Cell behaviour in Δ*sesB* mutant.** Lateral illumination was turned on at time 0 from the right side of the movie. The illumination was applied with three phases in the order of green, green/blue, and green with a time interval of 4 min. Δ*sesB* cell. Area 750 × 563 μm.

**Movie S11. Phototaxis of microcolony.** Lateral illumination of green and blue light was applied from the right side of the movie. WT cell at 45 °C. Area 117 × 88 μm.

**Movie S12. Phototaxis of cell perpendicular to the lateral light axis.** Lateral illumination of green and blue light was applied from the right side of the movie. WT cell at 45 °C. Area 171 × 44 μm.

**Movie S13. Phototaxis of cell that stand up and kept binding at a cell pole.** Lateral illumination of green and blue light was applied from the right side of the movie. WT cell at 45 °C. Area 171 × 44 μm.

**Movie S14. T4P dynamics through beads.** Extension and retraction of the T4P filament was visualized by sulfate beads with a size of 200 nm. WT cell at 45 °C. Area 42.0 × 31.5 μm.

**Movie S15. T4P dynamics through beads during negative phototaxis.** Lateral illumination was applied from the right side of the movie. Extension and retraction of the T4P filament was visualized by beads with a size of 200 nm. Dark-field microscopy. WT cell at 45 °C. Area 42.0 × 31.5 μm.
